# Building a Pangenome Alignment Index via Recursive Prefix-Free Parsing

**DOI:** 10.1101/2023.01.26.525723

**Authors:** Marco Oliva, Travis Gagie, Christina Boucher

**Affiliations:** Department of Computer and Information Science and Engineering, Herbert-Wertheim College of Engineering, Unversity of Florida, Gainesville, Florida, 32607, USA; Faculty of Computer Science, Dalhousie University, Halifax, Nova Scotia, Canada

## Abstract

**Motivation:** Pangenomics alignment has emerged as an opportunity to reduce bias in biomedical research. Traditionally, short read aligners—such as Bowtie and BWA—were used to index a single reference genome, which was then used to find approximate alignments of reads to that genome. Unfortunately, these methods can only index a small number of genomes due to the linear-memory requirement of the algorithms used to construct the index. Although there are a couple of emerging pangenome aligners that can index a larger number of genomes more algorithmic progress is needed to build an index for all available data.

**Results:** Emerging pangenomic methods include VG, Giraffe, and Moni, where the first two methods build an index a variation graph from the multiple alignment of the sequences, and Moni simply indexes all the sequences in a manner that takes the repetition of the sequences into account. Moni uses a preprocessing technique called *prefix-free parsing* to build a dictionary and parse from the input—these, in turn, are used to build the main run-length encoded BWT, and suffix array of the input. This is accomplished in linear space in the size of the dictionary and parse. Therein lies the open problem that we tackle in this paper. Although the dictionary scales nicely (sub-linear) with the size of the input, the parse becomes orders of magnitude larger than the dictionary. To scale the construction of Moni, we need to remove the parse from the construction of the RLBWT and suffix array. We accomplish this, in this paper by applying prefix-free parsing recursively on the parse. Although conceptually simple, this leads to an algorithmic challenge of constructing the RLBWT and suffix array without access to the parse. We solve this problem, implement it, and demonstrate that this improves the construction time by a factor of 8.9 the running time and by a factor of 2.7 the memory required.

**Availability:** Our implementation is open source and available at https://github.com/marco-oliva/r-pfbwt.

**Contact:** Marco Oliva at

## 1 Introduction

Read aligners—such as BWA [11] and Bowtie [10]—have been fundamental to the analysis of countless datasets, including the 1,000 Genome Project [19], the 100K Genome Project [20], the 1001 Arabidopsis Genomes project [21], and the Bird 10,000 Genomes (B10K) Project [14]. They have enabled the discovery of genetic markers that have causal relationships with countless diseases and phenotypes. These methods take as input a set of sequence reads and a reference genome, build an index from the reference genome, and use this index to find alignments with the limitation that few insertions and deletions are allowed.

Although short read aligners are sufficient for aligning to a single reference genome or a small number of reference genomes, they are unable to index the data of an entire population. To understand why, it is necessary to consider the underlying data structure used for constructing the index. The FM-index [5], which is a compressed subsequence index based on the Burrows Wheeler Transform (BWT) [4], is the primary data structure of the majority of these read aligners. The FM-index is not scalable to indexing thousands of genomes because it requires linear-space for its construction and storage. This reliance on the FM-index for read alignment is problematic as the research community has since recognized the necessity of pangenomics alignment, which aims to index and align to a population of genomes rather than a single genome. This has caused a systematic bias in biomedical research as the standard human reference genomes, such as the Genome Reference Consortium Human Build 38, come mainly from individuals of European ancestry. As Begley [1] states: *“*..*the reference genome falls short in ways that have become embarrassing, misleading, and, in the worst cases, emblematic of the white European dominance of science*..*”*

Currently, there exists a small handful of solutions for indexing and aligning to a pangenome, including Giraffe [18], VG [7], and Moni [17]. These methods follow different paradigms for pangenomics alignment; Giraffe [18] and VG [7] build and index a graph from a multiple alignment of the genomes. Moni [17] simply indexes all the reference genomes without any preprocessing, and takes advantage of the repetition in the reference genomes. Yet, even with these advancements, there still exists a critical gap in the number of genomes that are publicly available and our ability to index them for alignment. To understand our method, we give a brief overview of prefix-free parsing, which is the basis for our method.

Prefix-free parsing was first introduced by Boucher et al. [3] for building the BWT for a set of repetitive sequences, i.e., collection of genomes or chromosomes of the same species. The algorithm takes as input one or more set of strings (genomes) that are concatenated together using a delimiter symbol and two integers *w* and *p*. It then uses a rolling hash (such as the Karp-Rabin hash) to consider all *w*-length substrings and find all those that have hash values *h*, with *h* mod *p* = 0. These *trigger strings* are then uses to define a parse: each substring of the input string that begins and ends at a trigger string defines an string of the parse. All unique substrings of defined by the parse are stored in a dictionary, which is sorted lexicographically (alphabetically), and the parse can be stored in a compressed manner as the rank of the substrings in the dictionary. From the dictionary and parse, Boucher et al. [3] demonstrated that the BWT can be computed. Since then prefix-free parsing was extended to compute the suffix array (SA) [9], as a data structure [2] supporting multiple widely used query types, to construct a compressed suffix tree [15], to compress the input via Rpair [6], and to construct an an index for alignment of short reads [17].

## 2 Preliminaries

### 2.1 Basic definitions

A string *T* is a finite sequence of symbols *T* = *T* [1..*n*] = *T* [1] *⃛T* [*n*] over an alphabet Σ = *{c*_1_, …, *c*_*σ*_*}* whose symbols can be unambiguously ordered. We denote by *ε* the empty string, and the length of *T* as |*T* |. We denote as *c*^*k*^ as the string formed by the character *c* repeated *k* times.

We denote by *T* [*i*..*j*] the substring *T* [*i*] *⃛T* [*j*] of *T* starting in position *i* and ending in position *j*, with *T* [*i*..*j*] = *ε* if *i* > *j*. For a string *T* and 1 ≤ *i* ≤ *n, T* [1..*i*] is called the *i*-th prefix of *T*, and *T* [*i*..*n*] is called the *i*-th suffix of *T*. We call a prefix *T* [1..*i*] of *T* a *proper prefix* if 1 ≤ *i* < *n*. Similarly, we call a suffix *T* [*i*..*n*] of *T* a *proper suffix* if 1 < *i* ≤ *n*. Given a set of strings 𝒮, 𝒮 is *prefix-free* if no string in 𝒮 is a prefix of another string in 𝒮.

We denote by ≺ the lexicographic order: for two strings *T*_2_[1..*m*] and *T*_1_[1..*n*], *T*_2_ ≺ *T*_1_ if *T*_2_ is a proper prefix of *T*_1_, or there exists an index 1 ≤ *i* ≤ *n, m* such that *T*_2_[1..*i −* 1] = *T*_1_[1..*i −* 1] and *T*_2_[*i*] *< T*_1_[*i*]. Symmetrically we denote by ≺_colex_ the co-lexicographic order, defined to be the lexicographic order obtained by reading *T*_1_ and *T*_2_ from right to left instead that from left to right.

### 2.2 Suffix Array and Burrows-Wheeler Transform

Given a string *T* [1..*n*], the *suffix array* [13], denoted by SA_*T*_, is the permutation of *{*1, …, *n}* such that *T* [SA_*T*_ [*i*]..*n*] is the *i*-th lexicographically smallest suffix of *T*. We refer to SA_*T*_ as SA when it is clear from the context.

The Burrows-Wheeler transform of a string *T* [1..*n*], denoted by BWT_*T*_, is a reversible permutation of the characters in *T* [4]. If we assume *T* is terminated by a special symbol $ that is lexicographically smaller than any other symbol in Σ, we can define BWT_*T*_ [*i*] = *T* [SA_*T*_ [*i*]*−*1 mod *n*] for all *i* = 1, …, *n*.

### 2.3 Overview of Prefix-Free Parsing

Prefix-free parsing (PFP) takes as input a string *T* of length *n*, and two integers greater than 1, which we denote as *w* and *p*. It produces a parse of *T* consisting of overlapping phrases, where each unique phrase is stored in a dictionary. We denote the dictionary as D and the parse as P. We refer to prefix-free parse of *T* as PFP(*T*). As the name suggests, the parse produced by PFP has the property that none of the suffixes of length greater than *w* of the phrases in D is a prefix of any other. We formalize this property through the following lemma.

#### Lemma 1

([3]). *Given a string T and its prefix-free parse PFP* (*T*), *consisting of the dictionary* D *and the parse* P, *the set S of distinct proper phrase suffixes of length at least w of the phrases in* D *is a prefix-free set*.

The first step of PFP is to append *w* copies of # to *T*, where # is a special symbol lexicographically smaller than any element in Σ, and *T* does not contain *w* copies of #. For the sake of the explanation, we consider the string *T* ^*′*^ = #^*w*^*T* #^*w*1^. Next, we characterize the set of trigger strings E, which define the parse of *T*. Given a parameter *p*, we construct the set of trigger strings by computing the Karp-Rabin hash, *H*_*p*_(*t*), of substrings of length *w* by sliding a window of length *w* over *T* ^*′*^ = #^*w*^*T* #^*w*^, and letting E be the set of substrings *t* = *T* ^*′*^[*s*..*s*+*w−*1], where *H*_*p*_(*t*) *≡* 0 or *t* = #^*w*^. This set E will be used to parse #^*w*^*T* #^*w*^.

Next, we define the dictionary D of PFP. Given a string *T* and a set of trigger strings E, we let D = *{d*_1_, …, *d*_*m*_*}*, where for each *d*_*i*_ *∈ D*: *d*_*i*_ is a substring of #^*w*^*T* #^*w*^, exactly one proper prefix of *d*_*i*_ is contained in E, exactly one proper suffix of *d*_*i*_ is contained in E, and no other substring of *d*_*i*_ is contained E. Hence, we can build D by scanning #^*w*^*T* #^*w*^ to find all occurrences of the trigger strings in E, and adding to D each substring of #^*w*^*T* #^*w*^ that starts at the beginning of one occurrence of a trigger string and ends at the end of the next one. Lastly, the dictionary is sorted lexicographically. Given the sorted dictionary D and input string *T*, we can easily parse *T* into phrases from D with consecutive phrases overlapping by *w* characters. This defines the parse P as an array of indexes of the phrases in D. We note that *T* can then be reconstructed from D and P alone. We illustrate PFP using a small example. We let *w* = 2 and

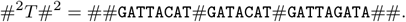

Now, we assume there exists a Karp-Rabin hash that define the set of trigger strings to be {AC, AG, T#, ##}. It follows that the dictionary D is equal to

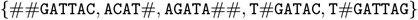

and the parse P to be [1, 2, 4, 2, 5, 3]. PFP can be used as a preprocessing step to build data structures such as the BWT and the SA.

In the next section, we will review how to compute the BWT of a string *T* using D and P.

### 2.4 Overview of Big-BWT

Given the prefix-free parsing of a string *T*, we show how to build the BWT of *T* using |*PFP* (*T*)| space, i.e., |D| + |P| space, where |D| is the sum of the length of its phrases and |P| the number of elements in it. This is referred to as the `Big-BWT` algorithm. We will use the following properties of PFP that were first introduced [3], and outlined in the following lemmas. Moreover, we will refer to them later when describing our recursive algorithm.

To determine the relative order of the characters in the BWT_*T*_ —and hence, the relative lexicographic order of the suffixes following those two characters in *T* —we start by considering the case in which two characters are followed in *T* by two distinct proper phrase suffixes *α, β ∈ S*. From Lemma 1 it follows

#### Corollary 1

([3]). *If two characters T* [*i*] *and T* [*j*] *are followed by different phrase suffixes α and β, where* |*α*| *≥ w and* |*β*| *≥ w, then T* [*i*] *precedes T* [*j*] *in the* BWT *of T if and only if α* ≺ *β*.

In other words, for some of the characters in BWT_*T*_, it is sufficient to only consider the proper phrase suffixes which follows them in *T* to break the ambiguity. When this is not enough, we need the information contained in the parse.

#### Lemma 2

([3]). *Let t and t*^*′*^ *be two suffixes of T that begin with the same proper phrase suffix α, and let q and q*^*′*^ *be the suffixes of P that have the last w characters of those occurrences of α and the remainders of t and t*^*′*^. *If t* ≺ *t*^*′*^ *then q* ≺ *q*^*′*^.

Next, we give some intuition on how these two lemmas are used to compute the BWT. For each distinct proper phrase suffix *α ∈ S* of length at least *w*, we store the range in the BWT containing the characters immediately preceding in the string *T* the occurrences of *α*. The starting position of the range for *α* is the sum of the frequencies in *T* (or P) of the proper phrase suffixes of length at least *w* that are lexicographically less than *α*. The length of the range is the frequency of *α*. Thus, we can store each proper phrase suffix along with their ranges in O(|D|) words of memory.

Now suppose that we are working on the range of the BWT of *T* corresponding to *i*-th proper phase suffix *α*. If all the occurrences of *α* in *T* are preceded by the same character `c`, then the range of the BWT of *T* associated with *α* will consist of all `c`’s. Therefore, no further computation is needed to define said range.

If the occurrences in the input of the *i*-th proper phrase suffix *α* are preceded by different characters then we make use of Lemma 2 to break the ambiguity. The order of the characters preceding *α* in *T* can be obtained from the order in which the phrases containing *α* appear in the BWT of P.

`Big-BWT` was later expanded to compute the SA values along with the BWT [9]. For each occurrence *α*_*i*_ of a proper phrase suffix *α ∈ S*, we can obtain its relative order among the other proper phrase suffixes using Lemma 2. In order to associate to *α*_*i*_ its SA value it is sufficient to store for each occurrence of each phrase *d*_*j*_ *∈* D its ending position in *T*. We denote the array containing the ending positions of *d*_*j*_ *∈* D as *EP*_*j*_. Let us assume that *α*_*i*_ is stored in the *k*-ht occurrence of the phrase *d*_*j*_ *∈* D, we are able to compute the associated SA value as *EP*_*j*_ [*k*] *−* |*α*_*i*_| + 1. Storing all the *EP* arrays requires O(|P|) words.

As described earlier, storing O(|P|) words can be a significant bottleneck for large repetitive datasets, and motivates the need for a BWT construction algorithm that uses less than O(|P|) words. In the next section, we introduce our recursive algorithm to address this need.

## 3 Methods

As previously mentioned, the size of D is at least one order of magnitude smaller than the size of P for large, repetitive input. This creates a computational bottleneck in building the *r*-index for pangenomes. We reduce this bottleneck by constructing the SA sample and RLBWT in memory that is significantly smaller than |P|+|D|. Previously, we removed this bottleneck in the construction of the RLBWT via recursive prefix-free parsing; however, construction of the SA samples together with the RLBWT proved to be more challenging. Here, we illustrate how both can be constructed via recursively prefix-free parsing, which resolves this open problem and makes construction of the *r*-index for pangenomes feasible. We begin by reviewing recursive prefix-free parsing, then we show how it can be extended to construct the SA samples at the same time as RLBWT.

### 3.1 Overview of Recursive Prefix-Free Parsing

We assume that PFP was ran on the input text *T* with window size *w*_1_ and integer *p*_1_. We denote the set of trigger strings defined by *w*_1_ and *p*_1_ as *E*_1_, and the output as P_1_ and D_1_. Next, we run PFP on the parse P_1_ with window size *w*_2_ and integer *p*_2_. We refer to running PFP on P_1_ as the *recursive step*, and denote the set of trigger strings defined by *w*_2_ and *p*_2_ as *E*_2_. We define the output of the recursive step as P_2_ and D_2_.

Next, we denote the set of proper phrase suffixes of length greater than *w*_1_ of the phrases in D_1_ as 𝒮 _1_ and, analogously, we denote the set of proper phrase suffixes of length greater than *w*_2_ of the phrases in D_2_ as 𝒮 _2_. Previously, we showed that BWT can be constructed in linear-time in |D_1_|, |D_2_|, and |P_2_|, which can be summarized as follows.

#### Theorem 1.

*[16] Given a string T, the dictionary* D_1_ *and the parse* P_1_ *obtained by running* PFP *on T, and the dictionary* D_2_ *and the parse* P_2_ *obtained by running* PFP *on* P_1_, *we can compute the* BWT_*T*_ *from* D_1_, D_2_ *and* P_2_ *using* O(|D_1_| + |D_2_| + |P_2_|) *workspace*.

We note that the important aspect about this Theorem is that it removes |*P*_1_| from the working space. Thus, we go from O(|D_1_| + |P_1_|) working space—where |*P*_2_| is quite large—to O(|D_1_| + |D_2_| + |P_2_|) working space. The proof of this theorem is constructive, meaning that we give a O(|D_1_| + |D_2_| + |P_2_|)-space construction algorithm to prove this result. Next, we briefly sketch this construction algorithm.

To define the construction algorithm, we note that—identical to `Big-BWT`—we consider each proper phrase suffix *α* in *S*_1_ and define its entries in the BWT. Here, we divide each proper phrase suffix into three possible subsets that we define: (1) easy proper phrase suffixes, (2) hard-easy proper phrase suffixes, and (3) hard-hard proper phrase suffixes. Figure 1 gives an illustration of these types. Based on the type of the proper phrase suffix, we calculate the RLBWT entry differently.

**Fig. 1:**
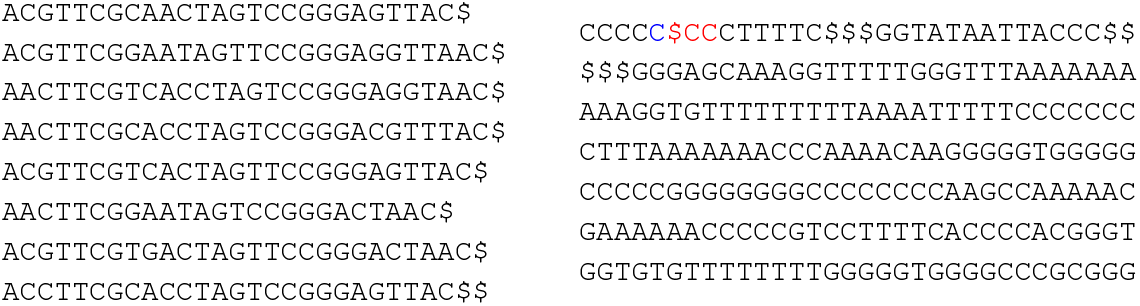
An illustration of a set of genomic sequences and their BWT. On the left is a set of genomic sequences that are concatenated together and delimited by `$`’s, with the last one delimited by `$$`. On the right is the BWT of the concatenated sequence. We will use the range BWT[4..8] as our ongoing example in this paper. The characters in the the range BWT[0..3] correspond to *easy* suffixes, the characters in blue to *hard-easy* suffixes, and the characters in red to *hard-hard* suffixes.

We begin by defining the first type of proper phrase suffix.

#### Definition 1.

*Given a proper phrase suffix α ∈ 𝒮* _1_, *stored in phrases D*_*α*_ = *{d*_*i*_, …, *d*_*k*_*}, with D*_*α*_ *⊂ D*_1_, *we denote α as an* easy *suffix if and only if in each d*_*j*_ *∈ D*_*α*_, *α is proceeded by the same character*.

As discussed in the Subsection 2.4, the ranges of the BWT of *T* corresponding to easy suffixes can be computed using only the information contained in D_1_. Hence, we can use `Big-BWT` for computing the BWT for these entries; no modification to the algorithm is needed. Next, we consider hard-easy proper phrase suffixes.

#### Definition 2.

*Given a proper phrase suffix α ∈ 𝒮* _1_, *stored in phrases D*_*α*_ = *{d*_*i*_, …, *d*_*k*_*}, with D*_*α*_ *⊂ D*_1_. *We let 𝒮* _*β*_ *⊂ 𝒮* _2_ *be the subset of proper phrase suffixes that are preceded by any of the phrases in D*_*α*_. *We denote α as an* hard-easy *suffix if and only if the following conditions are true: (a) there exists at least one pair of phrases* (*d*_*j*_, *d*_*l*_) *in D*_*α*_ *in which the character preceding α is different; and (b) no two elements of D*_*α*_ *precede in* P_1_ *the same proper phrase suffix in 𝒮* _*β*_ *⊂ 𝒮* _2_.

In other words, for hard-easy suffixes is enough to look at the proper phrase suffix in 𝒮 _2_ to break the ambiguity. We denote the remaining suffixes as hard-hard, which we define as follows.

#### Definition 3.

*Given a proper phrase suffix α ∈ 𝒮* _1_, *stored in phrases D*_*α*_ = *{d*_*i*_, …, *d*_*k*_*}, with D*_*α*_ *⊂ D*_1_. *We let 𝒮* _*β*_ *⊂ 𝒮* _2_ *be the subset of proper phrase suffixes that are preceded by any of the phrases in D*_*α*_. *We denote α as an* hard-easy *suffix if and only if the following conditions are true: (a) there exists at least one pair of phrases* (*d*_*j*_, *d*_*l*_) *in D*_*α*_ *in which the character preceding α is different; and (b) at least two elements of D*_*α*_ *precede in* P_1_ *the same proper phrase suffix in 𝒮* _*β*_ *⊂ 𝒮* _2_.

These definitions help us design a set of data structures to deal with the three different types of suffixes.

#### Definition 4.

*We define a table 𝒯* _1_ *containing* O(| *𝒮* _1_|) *rows and* O(1) *columns, such that for each α ∈ 𝒮* _1_, *we store in 𝒯* _1_ *its range in the* BWT *of T along with the co-lexicographic sub-range of the elements of* D_1_ *which store the occurrence of α. That is, for each c ∈* Σ_1_ *the columns of 𝒯* _1_ *store the range of co-lexicographically sorted phrases that end in α and have c in position* |*α*| + 1 *from the end*.

Next, we define the table 𝒯 _2_.

#### Definition 5.

*We define a table 𝒯* _2_ *containing* O(| *𝒮* _2_|) *rows and* O(1) *columns, such that for each α*^*′*^ *∈ 𝒮* _2_, *we store in 𝒯* _2_ *the co-lexicographic range of the phrases of* D_2_ *that contain α*^*′*^ *along with the meta-characters that precede α*^*′*^ *in P*_1_.

Lastly, we define the grid 𝒢 _2_.

#### Definition 6.

*We define the grid 𝒢* _2_ *containing* O(|*P*_2_|) *rows and* O(|*D*_2_|) *columns, such that for each element ℓ of* D_2_, *𝒢* _2_ *stores the positions in the* BWT *of* P_2_ *where ℓ appears*.

The data structures just defined allow us to compute the characters of the BWT of *T*. With 𝒯 _1_, we can compute all the ranges of the BWT of *T* corresponding to easy suffixes; for each easy suffix *α ∈ 𝒮* _1_ the corresponding range [*s*..*e*] of the BWT of *T* will be consisting of occurrences of the character, stored in 𝒯 _1_, preceding all the occurrences of *α* in *T*. We remind the reader that the extremes of that range can easily be computed from the number occurrences in P_1_ of each of the phrases in D_1_.

Using 𝒯 _2_, we are able to break the ambiguity in hard-easy suffixes, obtaining the relative order of the different characters preceding *α* in *T* as follows. We assume that the proper phrase suffix *α ∈ 𝒮* _1_ is preceded in *T* by multiple characters, making it a hard suffix. We iterate over the lexicographically sorted proper phrase suffixes of 𝒮 _2_ that are preceded in P_1_ by the phrases of *D*_1_ ending with *α*, and denote this set of phrases D_*α*_. We assume that a proper phrase suffix *β ∈ 𝒮* _2_ is preceded in P_2_ by only one element of D_*α*_, making the occurrence of *α*, i.e. *α*_*i*_, followed by *β*, a hard-easy suffix. Given that we are iterating over 𝒯 _2_ in lexicographic order, we know that *α*_*i*_ comes before all the following occurrences of *α*. Therefore, the BWT characters preceding *α*_*i*_ will come before the characters preceding the next occurrences of *α*. We now assume that while iterating over the proper phrase suffixes in 𝒮 _2_ that are preceded by the elements of D_*α*_, and are processing a proper phrase suffix *β ∈ 𝒮* _2_ that is preceded by more than one element of D_*α*_. In order to obtain the relative order of those occurrences of *β*, we consider the grid 𝒢 _2_ where each occurrence of the phrase storing *β* in the BWT of P_2_ will correspond to a character in the BWT of *T* in the order they appear in the BWT of P_2_.

Figure 2 and Figure 6 give an illustration of the two tables 𝒯 _1_ and 𝒯 _2_ we build for the genomes in Figure 1 and use to compute the BWT elements corresponding to easy and hard-easy suffixes respectively. We note that we put Figure 6 at the end of the paper due to its size. Figure 3 illustrates the grid 𝒢 _2_ used to compute the BWT elements corresponding to hard-hard suffixes.

**Fig. 2:**
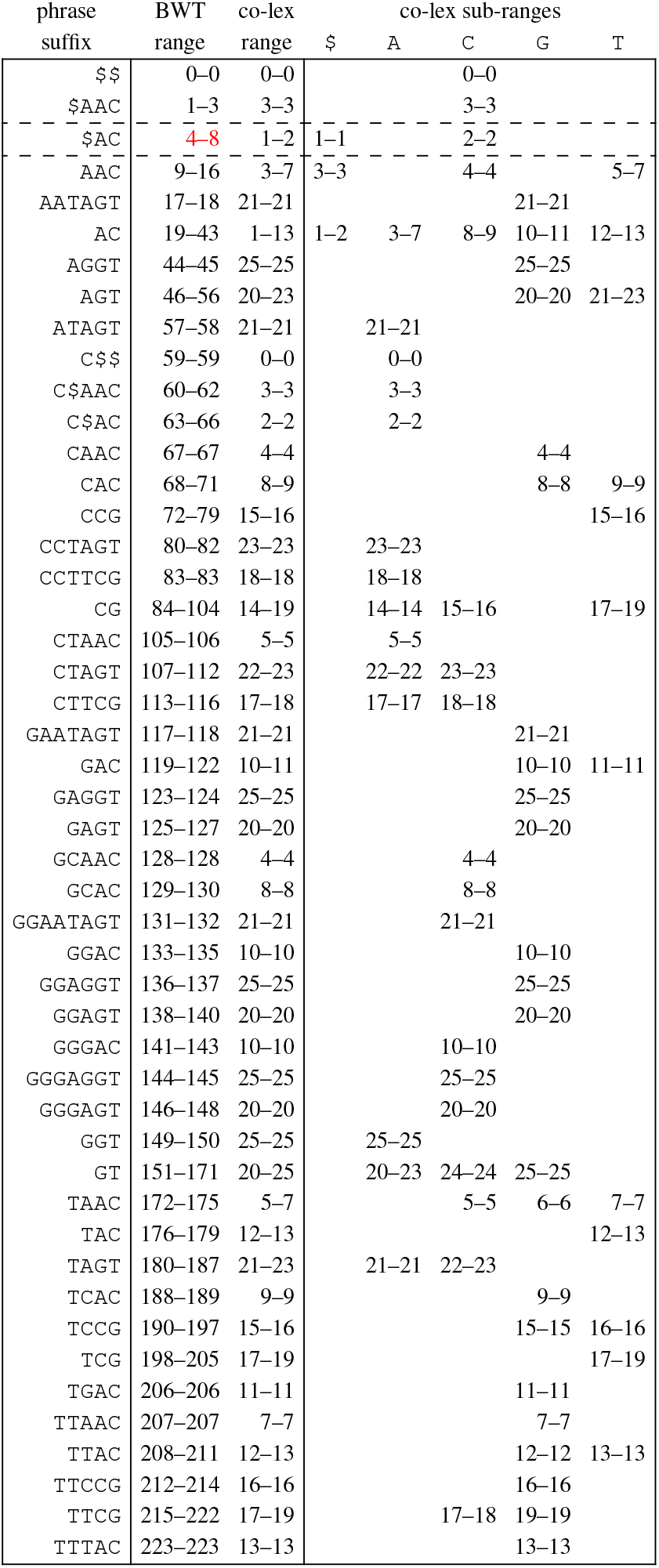
Our table 𝒯 _1_ storing all proper phrase suffixes of D_1_ of length at least *w*_1_ = 2.

**Fig. 3:**
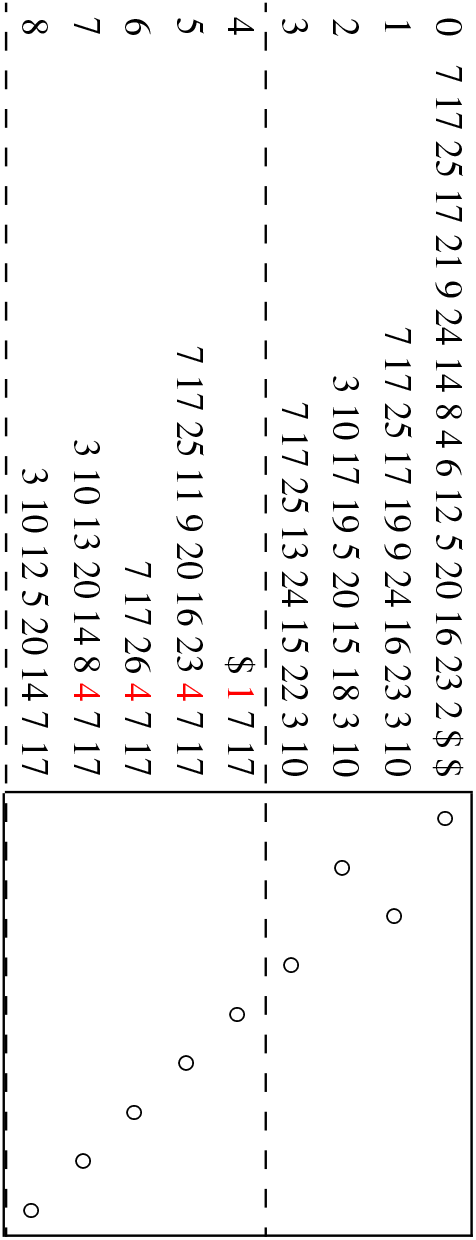
Our grid 𝒢 _2_ showing how the phrases (in co-lexicographic order of the expanded phrases) occur in the BWT of P_2_.

**Fig. 4:**
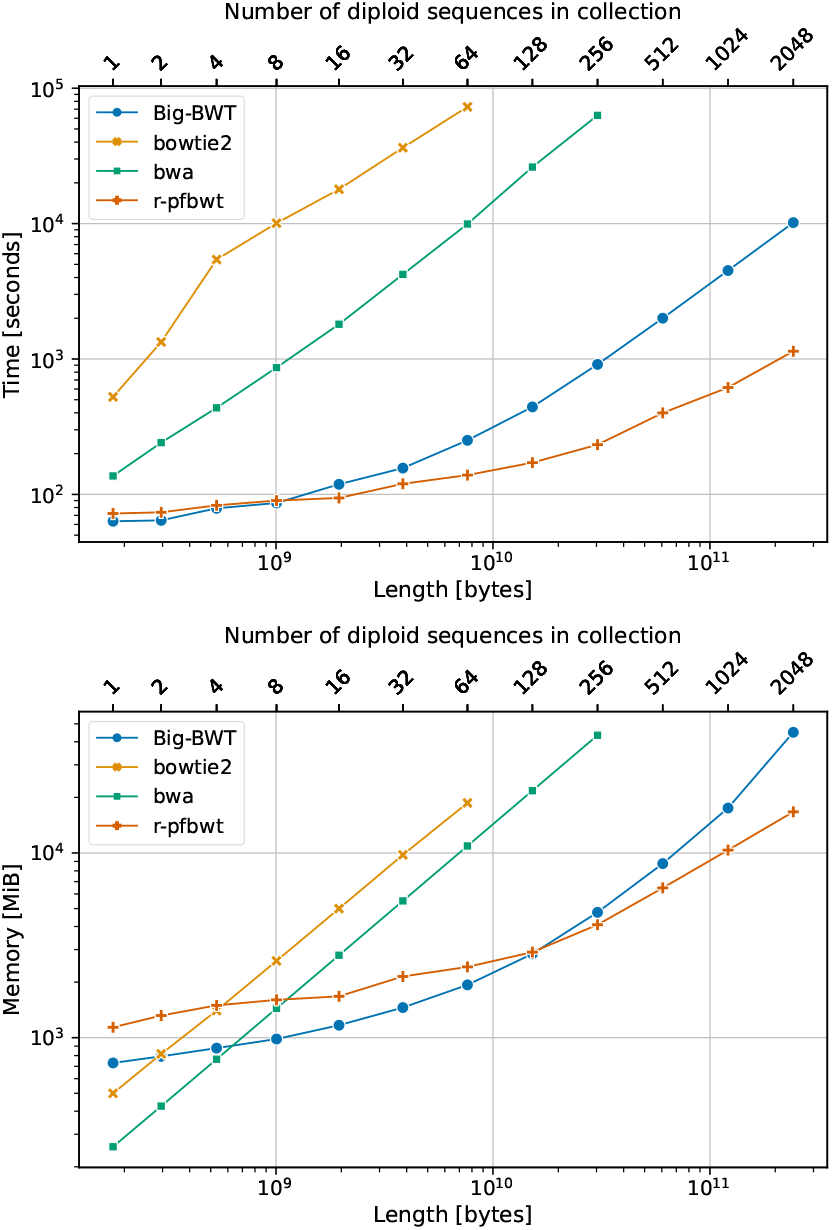
Chromosomes 19 construction Wall Clock Time in seconds (**top**) and peak memory in MiB (**bottom**)

**Fig. 5:**
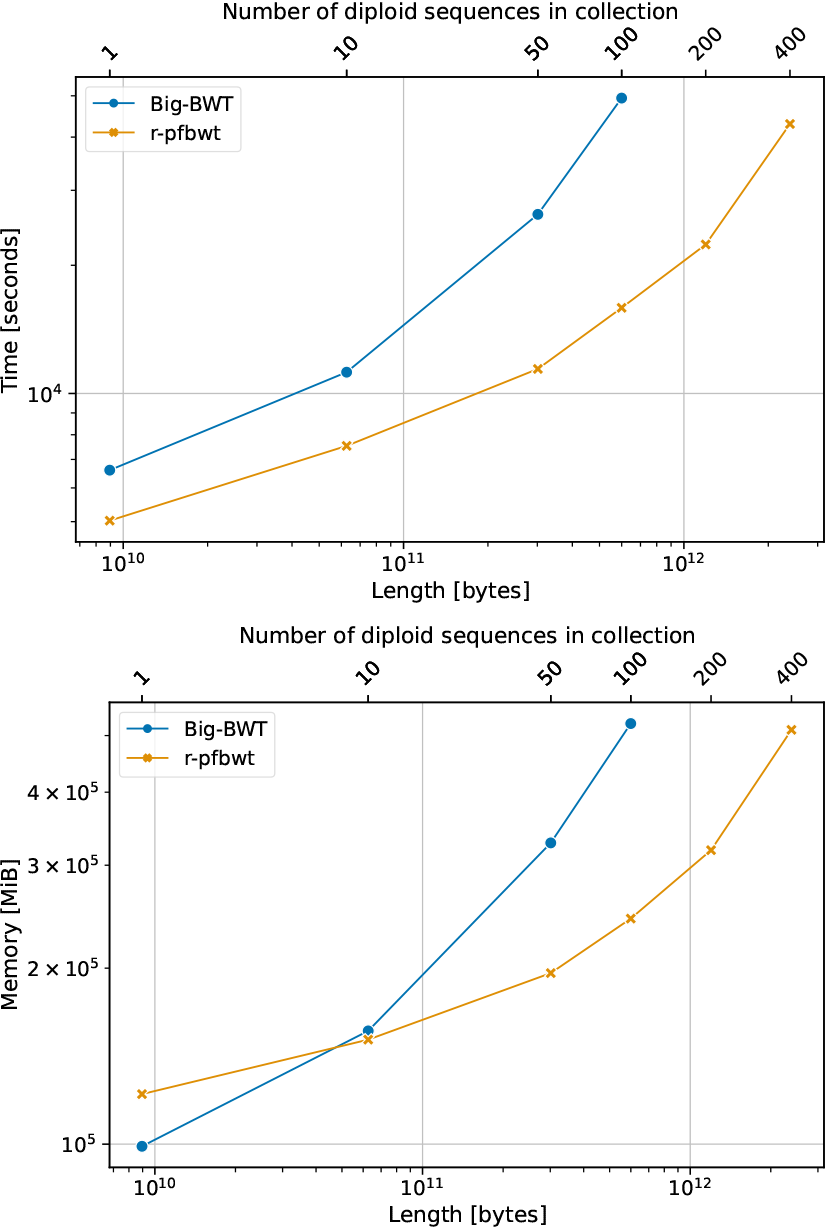
Human Genomes construction Wall Clock Time in seconds (**top**) and peak memory in MiB (**bottom**)

**Fig. 6:**
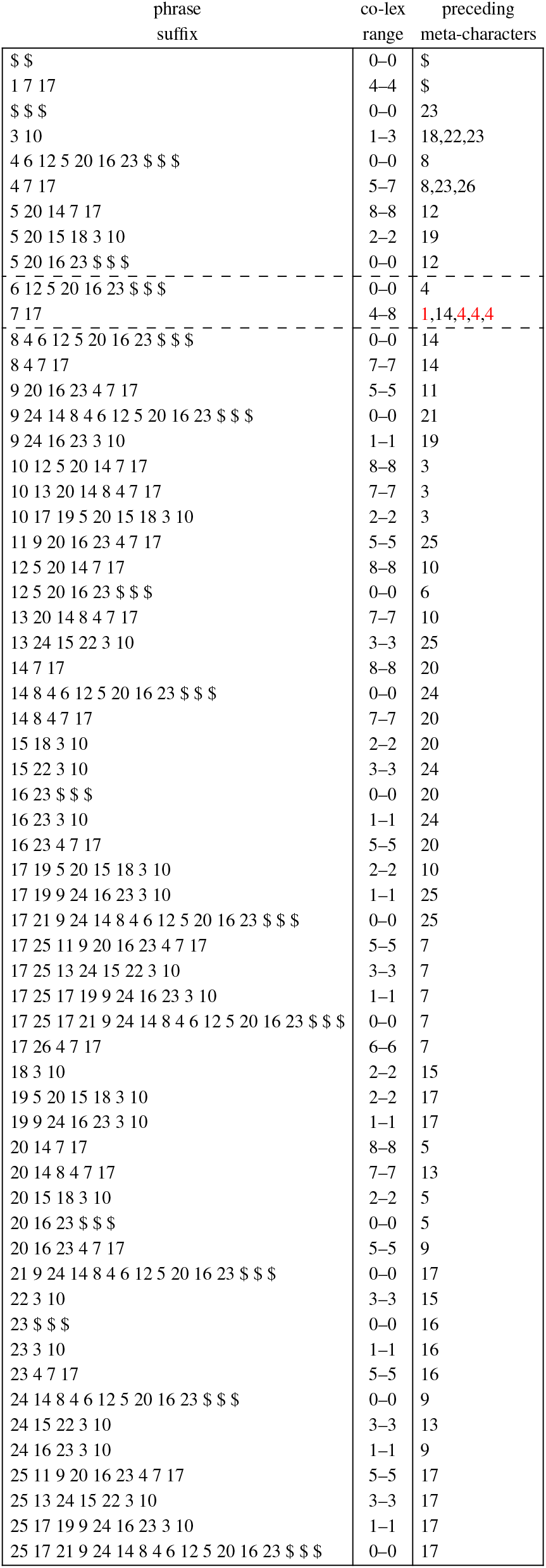
Our table 𝒯 _2_ storing all proper phrase suffixes of D_2_ of length at least *w*_2_ = 2.

To compute the BWT of the genomes in Figure 1 we iterate over the proper phrase suffixes of D_1_ stored in table 𝒯 _1_ in Figure 2. The first suffix is preceded in the input only by the character `C`, therefore the first suffix is an *easy* suffix and the associated range of the BWT (i.e. [0..0]) will consist of an occurrence of the character `C`. The same can be said for the second proper phrase suffix which defines the BWT range [1..3].

We now want to compute the range corresponding to the 3rd proper phrase suffix of D_1_ (i.e. `$AC`), namely the characters in blue and red in Figure 1.

From table 𝒯 _1_ in Figure 2, we know that `$AC` is preceded by `$`’s and `C`’s. For each phrase of D_1_ in the co-lexicographic range [1..2] (i.e. the two phrases `$$AC` and `AC$AC`, respectively encoded in P_1_ by the meta-characters `1` and `4`), we iterate over the proper phrase suffixes stored in 𝒯 _2_ preceded by those phrases in P_1_. In 𝒯 _1_, illustrated in Figure 6, we find that `6 12 5 20 16 23 $ $ $` is preceded in P_1_ by only the meta-character `4` making this occurrence of `$AC` a hard-easy suffix. We can extract the corresponding character from D_1_ from the phrase corresponding to the meta-character `4` (i.e. `C`).

We keep iterating over the proper phrase suffixes in 𝒮 _2_ that are preceded by either `1` or `4` and the next proper phrase suffix to consider is `7 17`, preceded by both the meta-characters `1` and `4`, making this occurrence of `$AC` a hard-hard suffix. To compute the corresponding BWT characters, we consider the occurrences in the BWT of P_2_ of the phrases encoding `7 17` in the co-lexicographic range [4..8]. From D_1_, we know that the occurrences of the phrase represented by the meta-character `4` will correspond to a `C` in the BWT while the occurrences of `1` to a dollar.

Following the order in which the phrases appear in the BWT of P_2_ stored in grid 𝒢 _2_ and illustrated in Figure 3 in the co-lexicographic range [4..8], we can define the range [6..8] of the BWT of the input as `$CC`.

### 3.2 Computing the Suffix Array values

In order to output the SA along with the BWT, we need to be able to compute the starting position in *T* of each occurrence of a proper phrase suffix *α ∈ 𝒮* _1_. The original solution proposed by Kunhle et al. [9] requires storing, for each occurrence in *T* of each phrase *d* in D_1_, its end positions in *T* in the array. We note that we use *α* to denote proper phrase suffixes and we use *d* to denote phrases in D_1_. We denote the array storing the ending position of each occurrence of *d ∈* D_1_ array as *EP*_*d*_. From the end position *EP*_*d*_[*j*] of an occurrence of *d* containing *α*_*i*_, we compute the start position of *α*_*i*_ by subtracting its length from *EP*_*d*_[*j*]. This strategy requires O(|P_1_|) words of memory. Here, we demonstrate that this can be accomplished only using D_2_ and P_2_; eliminating the need for P_1_. Even though in order to compute the BWT of *T* we introduced the three classes of proper phrase suffixes, we treat all the proper phrase suffixes as hard-hard. This is due to the fact that, when computing the range [*s*..*e*] of the BWT of *T*, if said range consists of copies of the same character *c*, we are not interested in the relative order of those occurrences. On the other hand, when computing the SA values the relative order matters to fully define the SA for the range [*s*..*e*]. Therefore, our solution relies on handling the relative order of the SA values as described for the hard-hard suffixes following from Lemma 3 [16].

Given that the phrases in D_2_ are on average short in practice (i.e. we set *p*_2_, which dictates the average phrase length, to be less than 15 in most cases), we can afford expanding the meta-characters of a phrase of *D*_2_ in order to compute the starting positions in *T* of an occurrence of a proper phrase suffix in *S*_1_. We formalize our approach as follows.

#### Theorem 2.

*Given a string T, the dictionary* D_1_ *and the parse* P_1_ *obtained by running* PFP *on T, and the dictionary* D_2_ *and the parse* P_2_ *obtained by running* PFP *on* P_1_, *we can compute the* SA *from* D_1_, D_2_ *and* P_2_ *using* O(|D_1_| + |D_2_| + |P_2_|) *workspace*.

Proof. We store for each occurrence in P_2_ of each element *ℓ ∈* D_2_ its starting position in *T* in an array *EP*_*ℓ*_. We let *t* = *T* [*i* + 1..*n*] be the suffix of *T* starting with *α*_*i*_ in phrase P_1_[*i*^*′*^]. Next, we let *q* be the suffix of P_1_ encoding the last *w*_1_ characters of *α*_*i*_ and the remainder of *t*. And we let *β*_*i*_ *∈ 𝒮* _2_ be the proper phrase suffix with which *q* starts. Next, we denote P_2_[*k*^*′*^] as the *j*-th occurrence of *ℓ*, encoding P_1_[*i*^*′*^] followed by *β*_*i*_. To simplify the notation, we denote |P_1_[*i*]| as the length of the phrase in D_1_ corresponding to the meta-character P_1_[*i*] (and analogously for |P_2_[*i*]|). Moreover, we denote |*ℓ*[*k*]| as the length of *d ∈* D_1_ with *d* = *ℓ*[*k*]. Given these definitions, we can compute the starting position of *α*_*i*_ in *T* as follows:

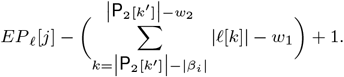

This computation only relies on P_2_, and can be accomplished for each *α*_*i*_ in *T*, leading to the SA. Hence, the SA can be computed along with the RLBWT in O(|D_1_| + |D_2_| + |P_2_|) workspace.

## 4 Experiments

We implemented `r-pfbwt` in ISO C++ 20. We used the `sdsl` library for `rank` and `select` support [8] and `gSACA-k` to compute the SA in our data structures [12]. `r-pfbwt` is open source and publicly available at https://github.com/marco-oliva/r-pfbwt. We designed two sets of experiments to evaluate the performance of `r-pfbwt`: (1) We compare `r-pfbwt` with Bowtie2 [10], BWA [11] and `Big-BWT` on 1, 2, 4, 8, 16, 32, 64, 128, 256, 512, 1024 and 2048 diploid sequences of chromosome 19 from the 1000 Genomes Project; (2) We select the methods that are able to successfully build their index on `chr19.2048` in less than 24 hours of Wall Clock Time and tested those methods on 1, 10, 50, 100, 200, 400 and 800 diploid human genomes sequences (excluding sex chromosomes) from the 1000 Genomes Project. Since Bowtie2 and BWA only take a FASTA file as input, we extracted the sequences from the VCF containing the variations and ran the methods on the extracted sequences. Both `Big-BWT` and `r-pfbwt`take as input the parse and dictionary produced by PFP. We remark that PFP can work directly from the VCF file without need to extract the sequences [15]. In reporting the results of both experiments, we do not take into account the resources needed to extract the sequences from the VCF file or to the resources needed to run PFP on the VCF file in order to focus the comparison on the index construction only. Both sets of experiments were ran on a node with AMD EPYC 7702 64-Core Processors for 128 cores and 1003 GB of memory with no other significant tasks running. All methods were ran with their default parameters.

### 4.1 Results on Chromosome 19

We ran Bowtie2, BWA, `Big-BWT` and `r-pfbwt` on exponentially increasing subsets of diploid sequences of Chromosome 19 from the 1000 Genomes Project. We generated the two haplotypes for each diploid sequence. Therefore, the number of DNA sequences in the set is twice the number of diploid sequences. We refer to the subsets as `chr19.1`, …, `chr19.2048`. Bowtie2 was able to build the index of `chr19.64` in 20 hours and 13 minutes, requiring 18GB of memory. Given that we set our limit to 24 hours, Bowtie2 was not able to build the index of `chr19.128`. BWA was able to build the index of `chr19.256` in 17 hours and 30 minutes requiring 43GB, and required more than 24 hours on `chr19.512`. Both `Big-BWT` and `r-pfbwt` were able to build the index for all the data points. On the largest input (i.e. `chr19.2048`) `r-pfbwt` was 8.9 times faster and required 2.7 less memory than `Big-BWT`. Yet, on the smaller data points, where the size of the D_1_ dominates, `Big-BWT` resulted faster and required less memory than `r-pfbwt`.

### 4.2 Results on Human Genome

Our second set of experiments aims to evaluate `r-pfbwt` and the competing methods on pangenome-scale datasets. We only considered `Big-BWT` and `r-pfbwt` for this experiment since these were the only methods capable of building their index on `chr19.2048`. We ran `Big-BWT` and `r-pfbwt` on increasingly larger subsets of the diploid human genomes sequences from the 1000 Genomes Project. We refer to the subsets as `hg.1`, `hg.10`, `hg.50`, `hg.100`, `hg.200` and `hg.400`. Given that `Big-BWT`’s original implementation is not able to handle a parse bigger than 16GB, for this set of experiments we used a newer version that fixes such problem at the expenses of a marginally longer execution time and bigger memory footprint. Both methods successfully build the index for `hg.1`, `hg.10` and `hg.50`. On `hg.10 r-pfbwt` required 2 hours and 5 minutes and 158GB of memory while `Big-BWT` required 3 hours and 6 minutes and 163GB. The difference between the two methods increased with he size of the dataset with `r-pfbwt` being 2.3 times faster and requiring 1.7 times less memory on `hg.50`. Furthermore, on `hg.100 r-pfbwt` was 3.1 times faster and required 2.2 times less memory. Lastly, `r-pfbwt` was able to complete the index construction on `hg.400` in 11 hours and 55 minutes requiring 536GB of memory.

## 5 Conclusion

In this paper, we presented a novel algorithm for removing the computational bottleneck created by the size of the parse in prefix-free parsing. In particular, we introduced an algorithm for building the SA sample and RLBWTof Moni in manner that removes the dependency of the construction on the parse from prefix-free parsing. This reduces the memory required by 2.7 times on large collections of chromosome 19. Moreover, neither Bowtie2 or BWA were able to index more than 256 copies of chromosome 19. On full human genomes this reducing was even more pronounced and `r-pfbwt` was the only method that was able to index 400 diploid human genomes sequences.

## Footnotes

1 We note that this definition is equivalent to original definition that considers the string *T* ^*′′*^ = *S*#^*w*^ to be circular.

## Notes

### Competing Interest Statement

The authors have declared no competing interest.

